# Computer Simulations Predict High Structural Heterogeneity of Functional State of NMDA Receptors

**DOI:** 10.1101/180091

**Authors:** Anton V. Sinitskiy, Vijay S. Pande

**Author notes:** Correspondence should be addressed to A.V.S or V.S.P.

## Abstract

It is unclear how the known atomic structures of neuronal NMDA receptors (NMDARs) relate to the functional states of NMDARs inferred from electrophysiological recordings. We address this problem by all-atom computer simulations, a method successfully applied in the past to much smaller biomolecules. Our simulations predict that four ‘non-active’ cryoEM structures of NMDARs rapidly interconvert on submicrosecond timescales, and therefore, correspond to the same functional state of the receptor.

## Introduction

N-methyl-D-aspartate (NMDA) receptors play the central role in the molecular mechanisms of higher cognitive abilities.^1,2^ Malfunctioning NMDA receptors (NMDARs) are involved in numerous disorders, including schizophrenia, epilepsy, intellectual disability, autism, Alzheimer’s, Huntington’s, Parkinson’s diseases, as well as neuronal damage following stroke.^3,4^ Recently, 18 X-ray and cryoEM structures of full-length NMDARs in various functional states (mainly closed and inhibited) have been reported.^5-9^ However, the dynamics of NMDARs at atomic resolution is poorly understood, as well as the correspondence of the published atomic structures to the dozens of the functional states of NMDARs inferred from electrophysiological recordings (Fig. S1).^2^

In this work, we modeled the structures of a ligand-free (henceforth referred to as ‘apo’) and an agonist-and-coagonist-bound (‘holo’) NMDAR at all-atom resolution (Fig. 1a, Video S1), and ran molecular dynamics (MD) simulations starting from these structures. To the best of our knowledge, these are the first reported simulations of a full NMDAR under physiological conditions. Previously, only parts of NMDARs,^10-16^ or an NMDAR without glycans and with a bound xenobiotic,^17^ were simulated. On the other hand, computer simulations of biomolecules at atomic resolution have been successfully employed in the past to study much smaller proteins (with hundreds of amino acid residues, vs. ∼3,000 residues in an NMDAR), resulting in detailed understanding of various metastable conformations and the dynamics of transitions between them.^18,19^ The rapid progress in hardware and software for MD simulations now enables us to extend this approach to neural receptors.

**Fig. 1.**
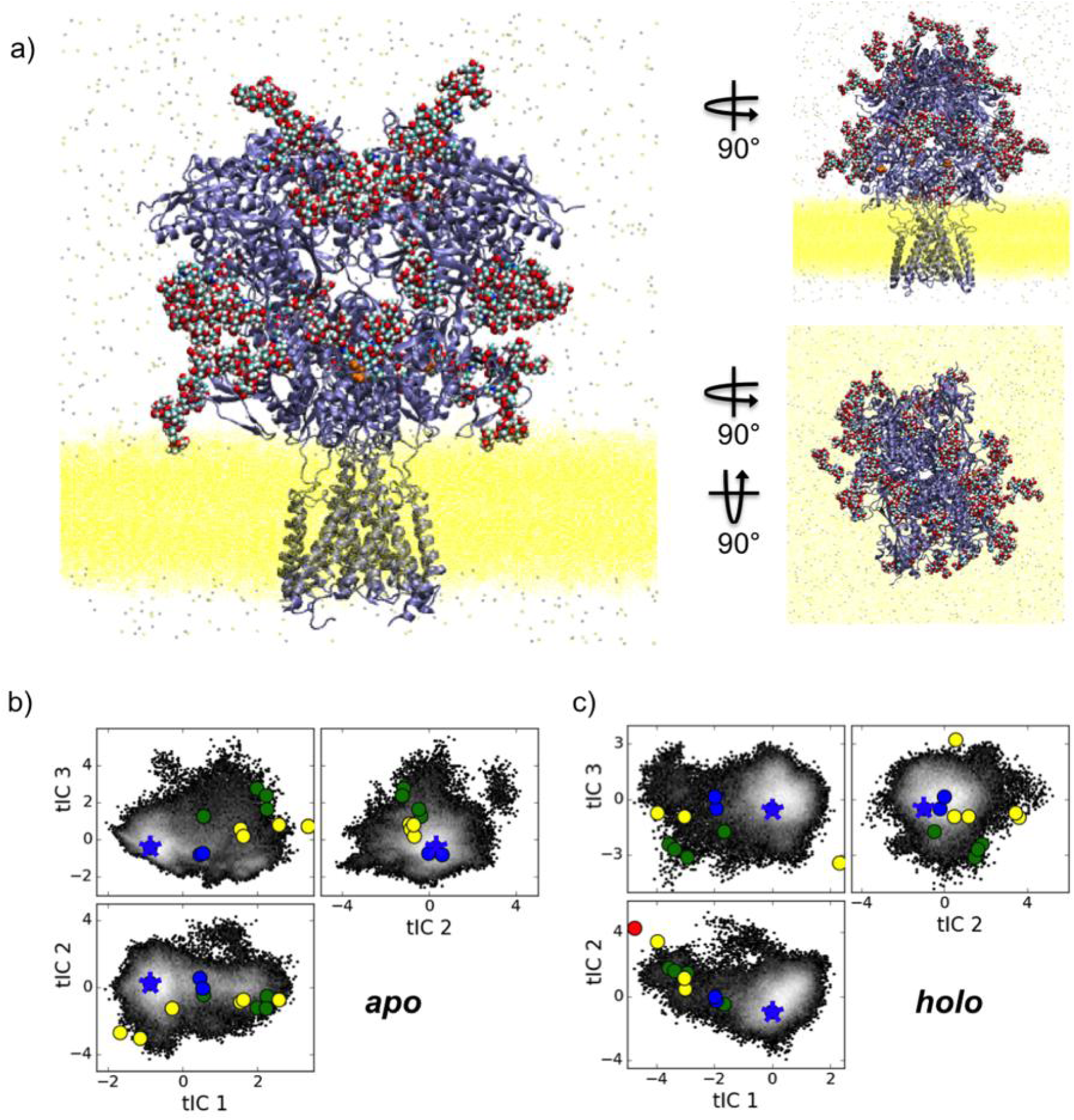
(a) We simulated a full NMDAR at all-atom resolution under physiological conditions, starting from the shown geometry. Protein part, *iceblue*; attached glycans, *vdW representation*; agonists (glutamate), *orange*; coagonists (D-serine), *orange*; lipid bilayer, *yellow*; NaCl, *gray*. The system was preequilibrated for 4.6 ns. (b,c) The range of NMDAR conformations sampled in our simulations included many geometries known from experiments. Conformational landscapes projected onto the three most informative time-structure based independent components (tICs) are shown. Computationally sampled conformations, *gray-to-black heatmap*; the initial geometry, *blue star*; structures of NMDAR in the inhibited state, *blue*; ‘non-active’ structures, *green*; ‘active’ structure, *red*; other structures, *yellow*.

## Results and Discussion

The conformations of an NMDAR generated in this work proved to be rather diverse, despite the fact that all MD trajectories started from the same geometry (Fig. 1b,c; Videos S2, S3). In particular, four ‘non-active’ cryoEM structures of NMDAR^7^ have all been reached in our simulations, though no information on these structures was used in the simulations in any way. One of the most striking differences between these four structures is in the degree of separation of the halves of the amino terminal domain (ATD) of the receptor from each other (Fig. 2). The distance between the ATD halves in the ‘non-active 2’ state^7^ (PDB code 5FXI) is ∼15 Å greater than in the ‘non-active 1’ state (5FXH), and ∼30 Å greater than in the inhibited state (4PE5), implying large-scale conformational transitions between these geometries. In our simulations, we have achieved an even greater diversity of structures, with the distance between the ATD halves ranging from 4 to 49 Å (Fig. 2a; Table S2). No transitions to the ‘active’ cryoEM structure^7^ or ‘classes 1-6’ found in antagonist-bound NMDARs^8^ occurred in the simulations (Fig. S6). On the other hand, some regions on the conformational landscape sampled in our simulations are not represented among the currently known experimental structures, in particular, the regions denoted as A-E in Fig. S9e and S10e (Section S6.3). Both of these results are reasonable, because the amount of sampling reached in this work could not suffice to observe transitions to all functional states of NMDARs, and the currently known structures are not expected to cover the whole variety of NMDAR conformations *in vivo*.

**Fig. 2.**
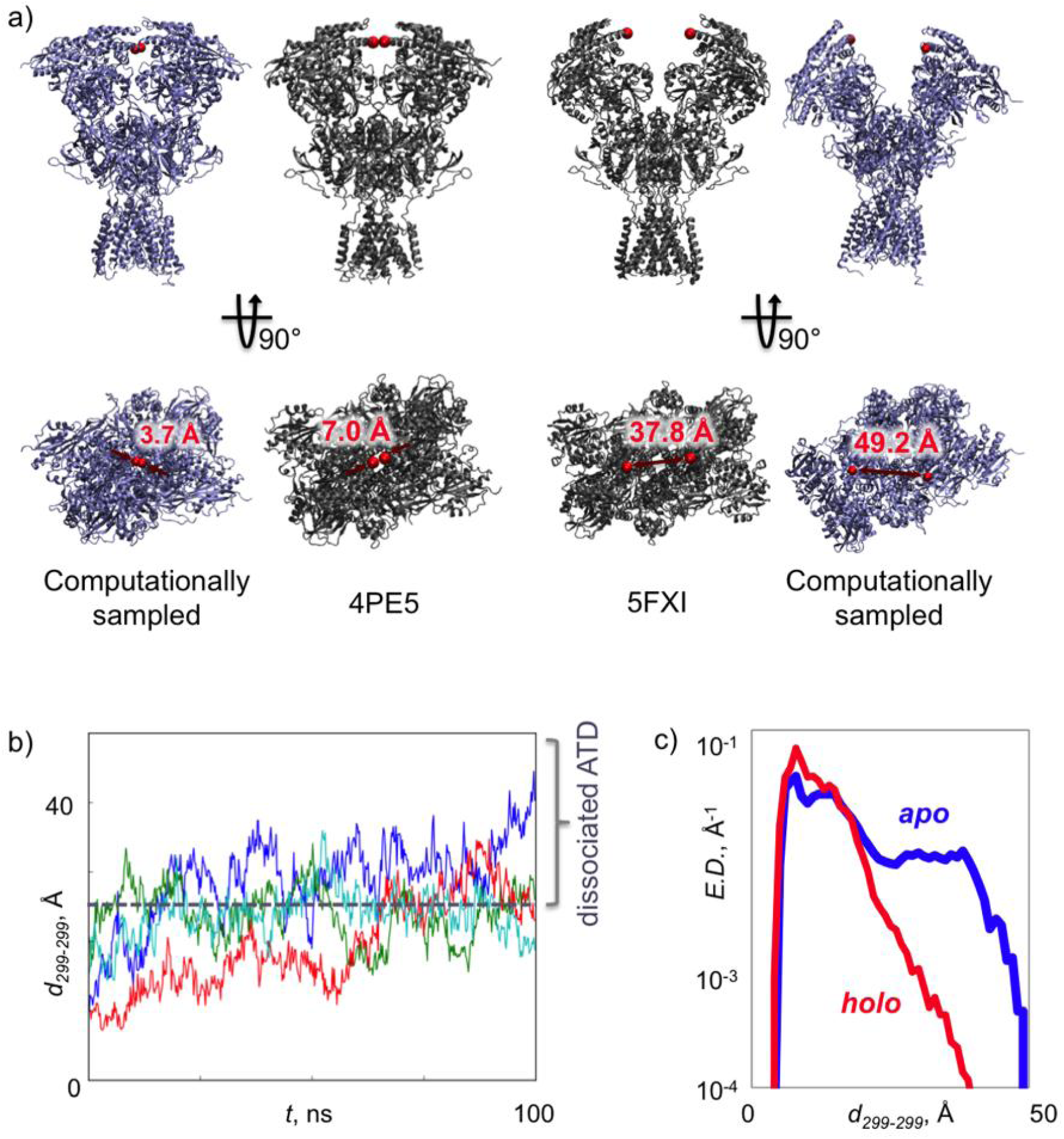
(a) The computationally sampled conformation with the smallest value of the distance *d*_*299-299*_ between two ATD halves has an even more tightly associated ATD than in the most tightly associated experimental structure, and the sampled conformation with the largest *d*_*299-299*_ is wider dissociated than many experimental structures (*only the protein part shown for clarity*). (b) Reversible dissociation of the ATD is directly visible from some of the MD trajectories. (c) Equilibrium distributions (E.D.) for the distance *d*_*299-299*_ in the holo (*red*) and apo (*blue*) NMDARs estimated with MSMs. Wider conformations are more stable in the apo state.

Timescales of conformational transitions between the sampled NMDAR geometries, including four ‘non-active’ conformations, can be up to hundreds of ns, as we predict here with the use of Markov state models (MSMs) based on the generated MD trajectories. This estimate is further corroborated by direct observations of a reversible dissociation of the ATD in a few MD trajectories over the timescale of less than 100 ns (Fig. 2b; Video S2). The fact that the predicted timescales are at least three orders of magnitude faster than the timescales for transitions between various functional states of NMDARs (Fig. S1b)^2,20^ implies that the sampled conformations, including four ‘non-active’ structures, refer to the same functional state of the receptor (specifically, one of the closed states).

Besides the timescales, MSMs can predict various equilibrium properties of the simulated system. In particular, we predict that the equilibrium fraction of more widely dissociated conformations is higher in the case of the apo than the holo form in the sampled functional state (Fig. 2c). Our simulations also reveal different dynamics in these two forms. Though the slowest degree of freedom (tIC 1, see Methods and Section S4) in both apo and holo forms turned out to be the same (a relative rotation of the ATD halves relative to each other in concert with a change in the distance between them), the second and third slowest degrees of freedom are different in these two forms (including a partial dilation of the gating ring in the apo form vs. more complicated conformational transitions in the holo form, see Section S4).

## Conclusions

To conclude, we approached a fundamental problem in understanding NMDA receptors, namely the problem of structures of NMDARs in various functional states, with unprecedented computer simulations of the full-length receptor at all-atom resolution. Our simulations demonstrate that large-scale transitions in the receptor (in particular, changes in the distance between the ATD halves by ∼40 Å) may occur astonishingly fast (on the submicrosecond timescale) and therefore, (a) functional assignment of X-ray and cryoEM structures cannot be based on the structural similarity alone and requires complementary information on the conformational dynamics, and (b) functional states of an NMDAR may each correspond to an ensemble of variegated conformations not reducible to a single geometry. More specifically, we conclude that four ‘non-active’ structures previously established by cryoEM^7^ refer to the same functional state (one of the closed states) of the receptor, based on the fact that their interconversions occur three orders of magnitude faster than transitions between different functional states known from electrophysiological experiments. Further work in this direction must carry out an experimental verification of the predictions of computer models. For the submicrosecond transitions predicted in this work, possible experimental techniques might be electron spin resonance and temperature-jump infrared spectroscopy. Computational prediction of new structures of NMDARs (Section S6.3) might also contribute to a better interpretation of cryoEM data.

## Methods

### Building the full all-atom structures of an NMDAR

A maximally complete structure of the protein part of an NMDAR was reconstructed based on all three X-ray structures available at the moment when the simulations started (PDB codes 4PE5,^5^ 4TLL, 4TLM^6^) (see Section S2). Mutations to the resulting structure were introduced to get the primary sequence as in a human GluN1/GluN2B NMDAR (uniprot codes Q05586-3, Q13224, http://www.uniprot.org). In the holo version, two glutamate molecules (agonist) were positioned as in the structure with PDB code 4PE5, and two D-serine molecules (coagonist) were added in positions corresponding to PBD structure 1PB8.^21^ Thirty four Man5 glycans were added to the receptor with the use of Glycoprotein Builder (http://glycam.org) to match the physiological glycosylation pattern.^22-25^ The glycosylated NMDAR was placed into a lipid bilayer with POPC and POPE (1:1) lipids and solvated in an explicit solvent (water) with a physiological concentration of NaCl of 0.154 M using CHARMM-GUI online tool (http://www.charmm-gui.org). The resulting systems contained 652,910 (apo) and 652,974 (holo) atoms.

Preequilibration was performed in Amber software^26^ separately for the apo and holo forms. The following force fields were used: ff99SB-ILDN^27^ for the protein part, GLYCAM_06i^28^ for the glycans, lipid14^29^ for the lipids, a force field by A. Horn^30^ for the zwitterion amino acids, and TIP3P^31^ for water. In each case, the potential energy was minimized over 10,000 steps (the first 5,000 steps with a harmonic restraint on all atoms in the protein part relative to the initial geometry, with the restraint factor of 1000 kcal/mol/Å^2^, and subsequent 5,000 steps without restraints). RMSD between the initial and energy-minimized geometries of the protein part was only 0.7 Å. After that, the system was gradually heated to 310 K over 4 ps in the NPV ensemble with a harmonic restraint on all atoms in the protein part relative to the geometry produced by the energy minimization, with the restraint constant of 100 kcal/mol/Å^2^. Further preequilibration of the system was performed in the NPT ensemble at 310 K and 1 atm in four steps: (1) with a harmonic restraint on all atoms in the protein part relative to the result of the previous stage of preequilibration, with the restraint factor of 10 kcal/mol/Å^2^, over 0.2 ns, (2) the same, with the restraint factor of 1 kcal/mol/Å^2^, over 0.4 ns, (3) the same, with the restraint factor of 0.1 kcal/mol/Å^2^, over 0.4 ns (4) without restraints, over 4 ns. RMSDs between the preequilibrated and the initial geometries were ∼2 Å both for the apo and holo forms. The preequilibrated geometries retained the secondary and tertiary structures present in the initial geometry. The difference between the initial and final geometries is mainly due to an overall swelling of the protein, which makes sense as the employed X-ray geometries were obtained at cryogenic temperatures, while the modeled structures refer to the body temperature (310 K).

### MD simulations

Production MD simulations were run in the NPT ensemble at 310 K and 1 atm with 2 fs timestep on graphical processing units (GPUs) in Amber software^26^ on Stanford computer cluster Sherlock. In total, 150 trajectories for the apo form and 150 trajectories for the holo form, each 100 ns long, have been generated. The performance of simulations depended on the type of GPUs automatically assigned on Sherlock, and reached ∼5.2 ns/day in the case of GeForce GTX Titan X GPU. The length of trajectories was limited to 100 ns because of two factors: first, lipid bilayer unphysically curved in ∼200-300 ns under the chosen simulation conditions, though such a distortion was not expected in the original paper for this lipid force field;^29^ second, the total amount of available compute resources was limited, and we chose to generate a greater number of parallel trajectories to have a more reliable statistics on the conformational transitions.

The simulations provide us with the coordinates of each of ∼600,000 atoms in the system recorded every 0.2 ns, which is an overwhelming amount of information. To enable a meaningful analysis, we processed these data in two stages: (1) featurization; (2) dimensionality reduction by time-structure based independent components analysis (tICA). More details on the data processing are given in Section S3.

### Featurization

Each frame from the MD trajectories was characterized by six distances, motivated by the previous analysis of experimental geometries of NMDARs^7^ (Fig. S3):

- *d*_*299-299*_ is defined as the distance between C_α_ atoms in residues 299 in two GluN1 subunits (in this work, the numeration of residues refers to the isoforms with uniprot codes Q05586-3 and Q13224). This variable quantifies the distance between the two heterodimers in the ATD.
- *d*_*178-184AB*_ and *d*_*178-184CD*_ are the distances between C_α_ atoms in residue 178 in a GluN1 subunit and residue 184 in a neighboring GluN2B subunit. They quantify the rotation of a GluN1 and a GluN2B subunit relative to each other within each of the two heterodimers.
- *d*_*247-247*_ is the distance between C_α_ atoms in residues 247 in two GluN2B subunits. It measures the rotation of the two heterodimers relative to each other in the ATD.
- *d*_*405-405*_ is the distance between C_α_ atoms in residues 405 in two GluN2B subunits. It quantifies a rotation of dimers relative to each other within the LBD, leading to changes in the distance between the ATD-LBD linkers.
- *d*_*658-658*_ is the distance between C_α_ atoms in residues 658 in two GluN2B subunits. It quantifies a rotation of dimers relative to each other within the LBD, leading to changes in the distance between the LBD-TMD linkers and, consequently, a dilation of the gating ring.

We did not include distances between two lobes (R1 and R2 or S1 and S2) in each subunit (GluN1, GluN2B) in the ATD and LTD domains into the list of the features, because we found it difficult to reduce the wealth of conformations of each subunit in each domain to changes in a small number of variables (data not shown). We will address the problem of the optimal choice of variables quantifying conformational transitions within these parts of NMDARs in the future work.

### tICA

Time-structure based independent components analysis (tICA)^32,33^ was used to automatically identify the slowest decorrelating degrees of freedom in the system, which are in most cases biomolecularly relevant. These degrees of freedom, called time-structure based independent components (tICs), are defined as the slowest-dynamics linear combinations of the input time series. The lagtime of 25 ns was used (significantly larger lagtimes would deteriorate the statistics required to compute temporal correlation functions and, therefore, tICs, while significantly shorter lagtimes might average out the slowest-timescale motions). tICA was performed on the six above-listed interatomic distances separately for the apo and holo form trajectories. In each case, the first three tICs were employed for further analysis.

The use of tICA allows for an automated selection of the most relevant degrees of freedom describing the modeled system. Like principal components in the widely used principal component analysis (PCA), time-structure based independent components (tICs) are linear combinations of the input data series. The difference between tICA and PCA is that tICs are defined as the degrees of freedom with the slowest decorrelating dynamics, while PCs are those with the widest scattering. tICs were shown to better capture biologically relevant degrees of freedom in proteins from MD simulations.^34^

A physical interpretation of the tICs computed for apo and holo NMDARs is provided in Section S4.

### MSMs

Markov state models were used to reconstruct equilibrium properties from finite-length MD trajectories and to estimate the timescales of the slowest conformational transitions in the system.^35^ To define the Markov states, the K-Centers clustering in the subspace of the first three tICs was performed. The number of clusters was chosen by cross-validation with the GMRQ technique.^36^ The optimal number of clusters turned out to be 197 and 79 for the apo and holo forms, respectively (Section S6.1). The lagtime for computing transition probabilities was taken to be 25 ns, the same as for tICA, in order to keep the balance between having sufficient statistics and retaining slowest motions. For the resulting probability transitions, MSMs were built separately for the apo and holo forms in the MSMBuilder software package (http://msmbuilder.org). An attempt to use the K-Medoids clustering instead of K-Centers led to MSMs with timescales of only up to tens of ns, thereby missing the slower conformational transitions in NMDARs successfully captured with K-Centers clustering. Assignment of the slowest timescales to specific transitions is given in Section S6.2, and a description of previously unknown structures of NMDARs predicted in this work is given in Section S6.3.

### Data availability

The data generated in this study, including the raw MD trajectories and the scripts used for the analysis, are publically available from Stanford Digital Repository by the following link: https://purl.stanford.edu/jf301dx7536.

## Acknowledgements

The authors were funded by NIH R01-GM062868. We thank Brooke Husic for valuable feedback on the manuscript.

## Author Contributions

A.V.S. conceived and designed the project, with supervision from V.S.P. A.V.S. carried out simulations, analyzed the results, wrote the paper and prepared supplementary materials. All co-authors interpreted the results. V.S.P. supervised the project and provided advice.

## Competing Financial Interests

V.S.P. is a consultant & SAB member of Schrodinger, LLC and Globavir, sits on the Board of Directors of Apeel Inc, Freenome Inc, Omada Health, Patient Ping, Rigetti Computing, and is a General Partner at Andreessen Horowitz. A.V.S declares no competing financial interests.

## SUPPLEMENTARY INFORMATION

### S1. Structural vs. functional data on NMDA receptors

In the literature, there is a wide gap between functional models of NMDA receptors (Fig. S1a,b) and their known atomically resolved structures (Fig. S1c). Though most experimentally determined structures, except for the structure with PDB code 5FXG, are believed to refer to functional closed states, it is not clear which of the closed states shown in Fig. S1a,b each of the structures belongs to, and what degree of structural diversity is typical of each functional state. We address this problem in the present paper with atomically resolved computer simulations of NMDA receptors.

**Fig. S1.**
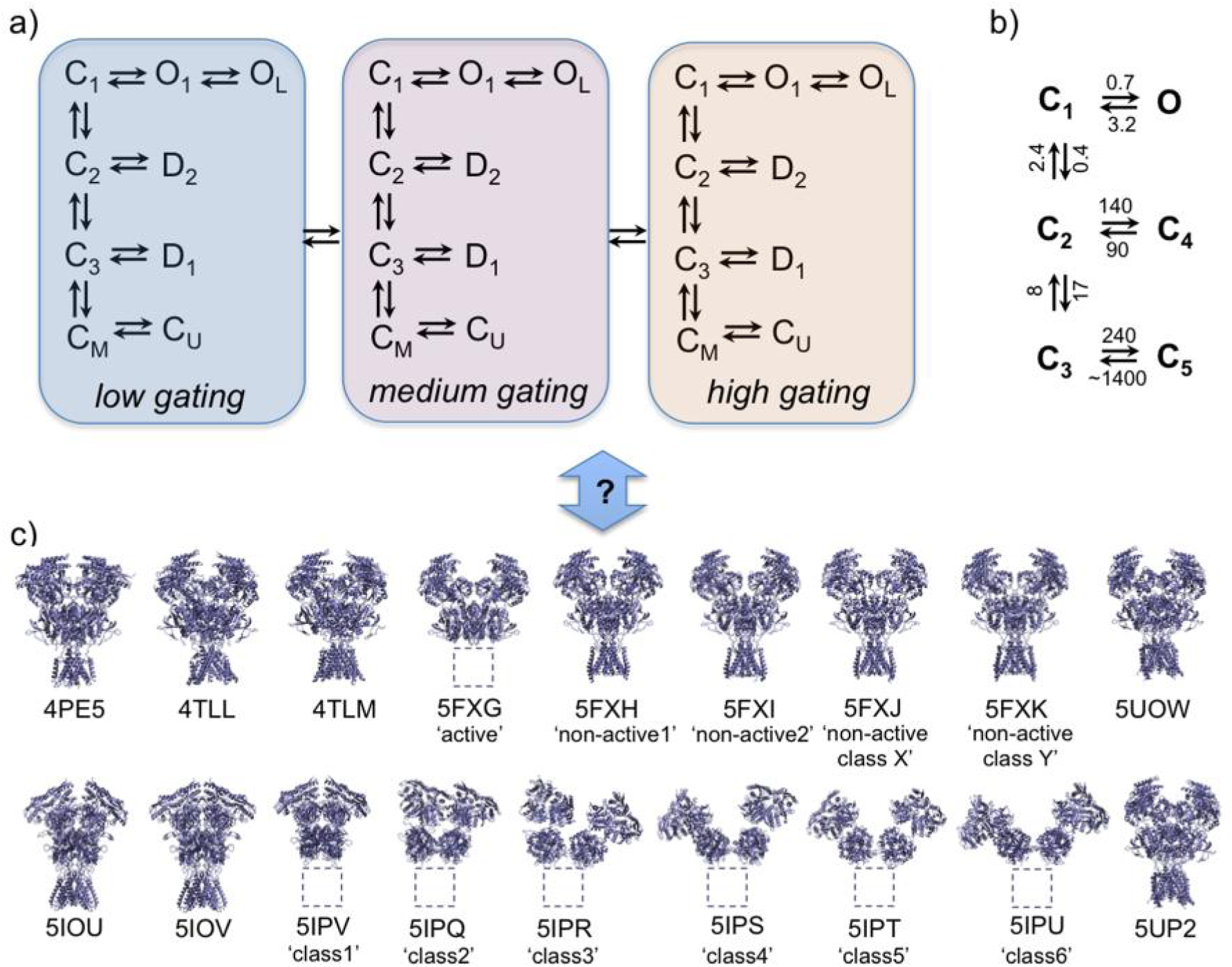
Multiple functional states of NMDARs were inferred from electrophysiological experiments. (a) A tiered model of an NMDAR includes 27 states. (b) A simplified model includes 5 closed and one open states. Characteristic timescales (defined as inverse rate constants) are given in ms. (c) The correspondence between these functional states and 18 atomic structures of NMDARs is unclear.

### S2. Building the all-atom structures of a full-length NMDAR

By the time we started the simulations, only three structures of the full-length NMDA receptor were available (PDB codes 4PE5,^5^ 4TLL, 4TLM^6^). Aiming at modeling a human NMDAR, we chose the structure with PDB code 4PE5 as the main source of information. This structure is for an NMDAR from a mouse (*Rattus norvegicus*) and differs from the human NMDAR only by point mutations. The structure with PDB code 4PE5 contains 90% of the residues in the extracellular and transmembrane domains (including residues with point mutations). The other two structures (PDB codes 4TLL and 4TLM) refer to a frog (*Xenopus laevis*) and differ from the human NMDAR not only by point mutations, but also insertions and deletions. We used these two structures to fill out gaps in the 4PE5 structure. As a result, we reconstructed the geometry of most of the extracellular and transmembrane parts of the human NMDAR (residues 23-584, 601-843 in the GluN1 subunit and residues 30-578, 599-845 in the GluN2B subunit). The resulting structure is missing the initial, presumably highly disordered, fragments of both subunits in the amino terminal domain (ATD) (residues 1-22 in GluN1 and 1-29 in GluN2B) and the intracellular C-terminal domain (CTD) (residues 585-600, 844-938 in GluN1 and 579-598, 846-1484 in GluN2B). All these fragments are missing in all currently available experimental structures of NMDARs, despite that they are usually referred to as full-length structures.^5-9^

We used the following algorithm to model the full-length structure of an NMDAR:

1. Wherever possible, we filled out gaps in the 4PE5 structure by inserting fragments available from the other chain in the same structure (chains A and C both correspond to GluN1 subunits and are related by an approximate C_2_ symmetry, and the same holds for chains B and D corresponding to GluN2B subunits). Before insertion, the corresponding fragments were manually aligned with the neighboring parts of the protein backbone present in the original 4PE5 structure:
  - if two residues on each side from the missing fragment were present, we aligned the fragment to be inserted by two resides on each side (here and below, the alignment was performed by RMSD for the protein backbone),
  - if two residues were present only on one side of the missing fragments, and the other side was not terminal, we aligned the fragment to be inserted by two residues on the side where the neighboring residues were present, and one closest residue on the side where the immediate neighbors were not resolved.
  - for terminal residues, we aligned the inserted fragment by the three residues on the side with a resolved structure.
2. After that, we filled out the remaining gaps in the protein backbone with fragments from 4TLL and 4TLM structures. When more than one variant of the missing fragment were available, we chose one with the smallest RMSD computed after alignment.
3. Residues 544 and 545 in the GluN2B subunit were missing in all three experimental structures, and could not be reconstructed at steps 1 and 2. We manually added two lines to the structure file for the C_α_ atoms in these two residues in each chains. The coordinates of these C_α_ atoms were computed by a linear extrapolation between the coordinates of the C_α_ atoms in residues 543 and 546. The resulting local errors in the structure were eliminated at the stage of structure minimization and preequilibration.
4. To get the primary sequence of the protein corresponding to the human NMDAR, we mutated the residues whenever necessary by manually editing the structure file (replacing the name of the amino acids and deleting the lines for the atoms that are not in common between the original and final amino acids).

The fragments inserted into the modeled structure according to this algorithm are listed in Table S1 and visualized in Fig. S2.

**Table S1.** Fragments of the protein inserted into the modeled structure.

| **chain** | **residues missing in 4PE5** | **origin of the inserted fragment** | **resides used for alignment** |
| --- | --- | --- | --- |
| **A** | 23-24 | 4TLM, chain A | 25-27 |
|  | 53-57 | 4PE5, chain C | 51, 52, 58, 59 |
|  | 95-102 | 4TLL, chain A | 93, 94, 103, 104 |
|  | 442-444 | 4TLM, chain A | 440, 441, 445, 446 |
|  | 445-449 | 4PE5, chain C | 441, 450, 451 |
|  | 583-584 | 4TLM, chain C | 581, 582, 605 |
|  | 601-604 | 4TLM, chain A | 582, 605, 606 |
|  | 617-622 | 4TLM, chain C | 615, 616, 623, 624 |
|  | 657-663 | 4PE5, chain C | 655, 656, 664, 665 |
|  | 802-808 | 4PE5, chain C | 800, 801, 809, 810 |
|  | 831-833 | 4PE5, chain C | 828-830 |
|  | 834-843 | 4TLM, chain A | 831, to, 833 |
| **C** | 23-24 | 4TLM, chain A | 25-27 |
|  | 95-102 | 4TLL, chain A | 93, 94, 103, 104 |
|  | 442-444 | 4TLM, chain A | 440, 441, 445, 446 |
|  | 549-553 | 4PE5, chain A | 547, 548, 554, 555 |
|  | 583-584 | 4TLM, chain A | 581, 582, 605 |
|  | 601-604 | 4TLM, chain A | 582, 605, 606 |
|  | 617-622 | 4TLM, chain C | 615, 616, 623, 624 |
|  | 623-633 | 4PE5, chain A | 616, 634, 635 |
|  | 834-843 | 4TLM, chain A | 831-833 |
| **B** | 395-397 | 4TLL, chain D | 393, 394, 404 |
|  | 398-402 | 4TLM, chain B | 394, 403, 404 |
|  | 440-450 | 4TLL, chain D | 438, 439, 451, 452 |
|  | 540 | 4PE5, chain D | 538, 539, 551 |
|  | 541-543 | 4TLM, chain D | 539, 540, 550 |
|  | 546-549 | 4TLL, chain B | 540, 550, 551 |
|  | 550 | 4PE5, chain D | 539, 551, 552 |
|  | 570-578 | 4TLM, chain B | 568, 569, 602 |
|  | 599-601 | 4TLL, chain D | 569, 602, 603 |
|  | 616-629 | 4TLM, chain B | 614, 615, 630, 631 |
|  | 803-806 | 4TLL, chain D | 801, 802, 807, 808 |
|  | 807-820 | 4PE5, chain D | 802, 821, 822 |
|  | 842-845 | 4TLM, chain D | 839, 840, 841 |
| **D** | 30-31 | 4PE5, chain B | 32-34 |
|  | 395-397 | 4TLL, chain D | 393, 394, 404 |
|  | 398-402 | 4TLM, chain B | 394, 403, 404 |
|  | 440-450 | 4TLL, chain D | 438, 439, 451, 452 |
|  | 451 | 4PE5, chain B | 439, 452, 453 |
|  | 541-543 | 4TLM, chain D | 539, 540, 550 |
|  | 546-549 | 4TLL, chain B | 540, 550, 551 |
|  | 568-569 | 4PE5, chain B | 566, 567, 602 |
|  | 570-578 | 4TLM, chain B | 568, 569, 602 |
|  | 599-601 | 4TLL, chain D | 569, 602, 603 |
|  | 615 | 4PE5, chain B | 613, 614, 630 |
|  | 616-629 | 4TLM, chain B | 614, 615, 630, 631 |
|  | 803-806 | 4TLL, chain D | 801, 802, 807, 808 |
|  | 842-845 | 4TLM, chain D | 839, 840, 841 |
|  | 839-841 | 4PE5, chain B | 836-838 |

All missing atoms, both absent in the experimental structures and deleted during the structure modeling at the previous stages, were added back with the use of *tleap* tool from AmberTools software package.^26^

Thirty four glycans were added to the receptor to match the physiological glycosylation pattern (N-glycosylated residues 61, 203, 239, 276, 300, 350, 368, 440, 471, 491, 771 in the GluN1 subunit and 74, 341, 348, 444, 491, 688 in the GluN2B subunit).

The glycosylated NMDAR was placed into a lipid bilayer. The upper leaflet of the bilayer contained 248 molecules of POPC and 248 molecules of POPE, and the lower leaflet contained 243 molecules of each.

Having started this sentence with a letter rather than a digit, 156,365 water molecules, 469 Na^+^ ions and 407 Cl^-^ ions were added to neutralize the system and ensure the physiological level of the ionic strength of the solution surrounding the NMDAR. The resulting unit cell had the dimensions of 189 × 189 × 215 Å.

To remove steric clashes and other issues in the modeled structure, the system was preequilibrated as described in Online Methods.

**Fig. S2.**
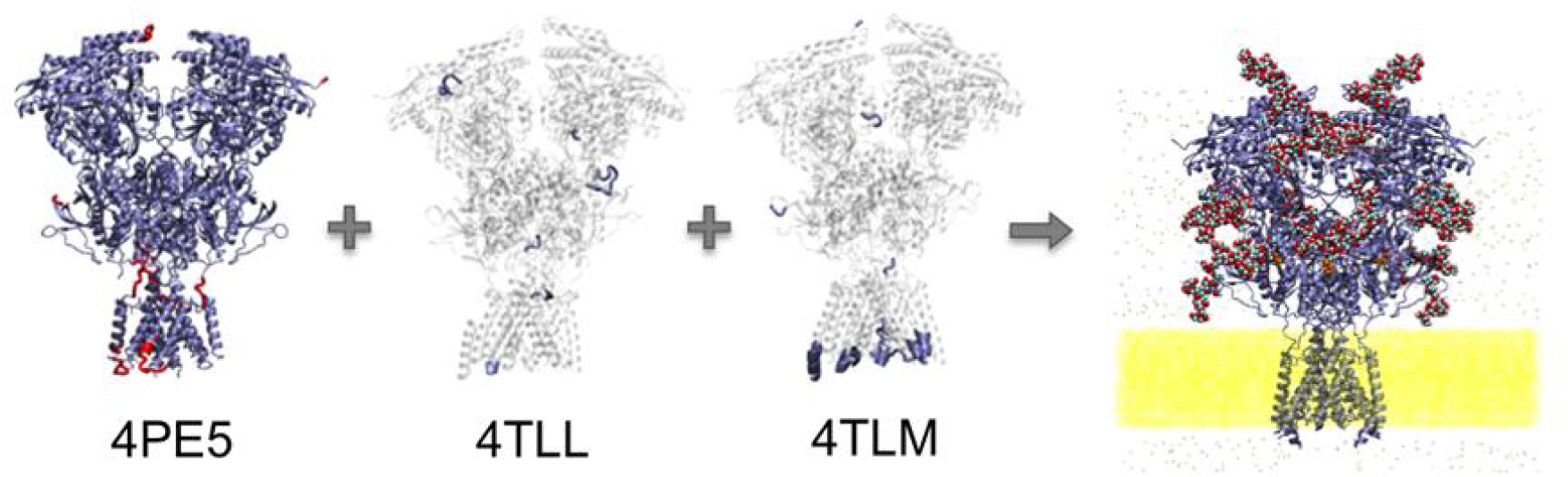
The X-ray structures of the receptor with PDB codes 4PE5 (predominantly), 4TLL and 4TLM were used to reconstruct the structure of the protein part of the full-length NMDAR. Used parts of the X-ray structures are shown in blue; parts of 4PE5 used twice, for the same and the other symmetric chains, are shown in red.

Having finished the production simulations, it has not escaped our notice^37^ that the side chain of residue Val618 of chain D unphysically penetrated into the ring in residue Trp607 of the same chain in the modeled structure. This steric issue has not been resolved by the energy minimization and preequilibration, and retained in all produced MD frames. We believe that this issue has not affected the conclusions made in this paper, because its physical effect is an additional stabilization of the secondary structure of one of the helices in the transmembrane domain, which is by itself a stable fragment. Our simulations sampled conformational transitions only in the extracellular part of the receptor, while the transmembrane domain stayed stable and closed in all the trajectories, and therefore, this steric issue should not have had a noticeable effect on the results.

The question of where the lipid bilayer lies relative to NMDAR and AMPA receptors has not been directly addressed so far, though the existing notion about it can be seen from several cartoon representations given in previously published papers.^1,8,9,38,39^ In our simulations, the receptor submerged into the lipid bilayer deeper than expected, and some glycans and amino acid residues from the extracellular part of the NMDAR interacted with lipids in the simulated structures. These interactions might affect the structure of the receptor by widening the lower part of the LBD and slowing down conformational transitions in the LBD due to glycans serving as anchors stuck in the lipid bilayer. Based on our simulations, we formulate a hypothesis for further experimental investigation that glycans on NMDARs might directly interact with lipids from the cell membrane.

### S3. Data processing

#### S3.1. Damaged data omission

In 3 of 150 MD trajectories for the apo NMDAR, individual frames were omitted from trajectories used for the analysis, because these three frames were damaged in the middle of output for unknown reasons.

- In trajectory 42, the frame corresponding to *t* = 32.0 ns was omitted. Therefore, the resulting 100 ns trajectory used for further analysis actually consists of two concatenated fragments corresponding to *t* = 0.2 … 31.8 ns and *t* = 32.2 … 100.2 ns in the simulation.
- In trajectory 96, one frame corresponding to *t* = 39.0 ns was omitted.
- In trajectory 130, one frame corresponding to *t* = 30.2 ns was omitted.

Since the time shift in each of these three trajectories caused by omitting a frame (0.2 ns) is much less than the lag times used for computing tICs and building MSMs (25 ns), and since there are no noticeable conformation changes in the vicinities of all three omitted frames, we believe that omitting these three frames did not affect in any way the reported results.

The other 147 trajectories for the apo NMDAR and all 150 trajectories for the agonist- and-antagonist-bound NMDAR do not have missing fragments and correspond to *t* = 0.2 … 100.0 ns in the corresponding simulations.

#### S3.2. Featurization

For each frame in the simulated MD trajectories, six distances were computed as described in Online Methods (see also Fig. S3). These distances were defined in such a way that they capture the main conformational changes in the NMDAR discussed in the previous literature on the analysis of the experimental structures of NMDARs.^7,8^ In general, computer modeling can reveal previously unknown biologically relevant degrees of freedom in simulated systems.^18,19,33,34^ However, this type of investigation requires more sampling than an analysis based on the known collective variables, which is currently difficult to achieve for such a large system as a full-length NMDAR. For this reason, in the present work we limit ourselves to an analysis based on these six interatomic distances motivated by the previous literature, leaving the question of an automatic detection of all biomolecularly relevant degrees of freedom for future work.

**Fig. S3.**
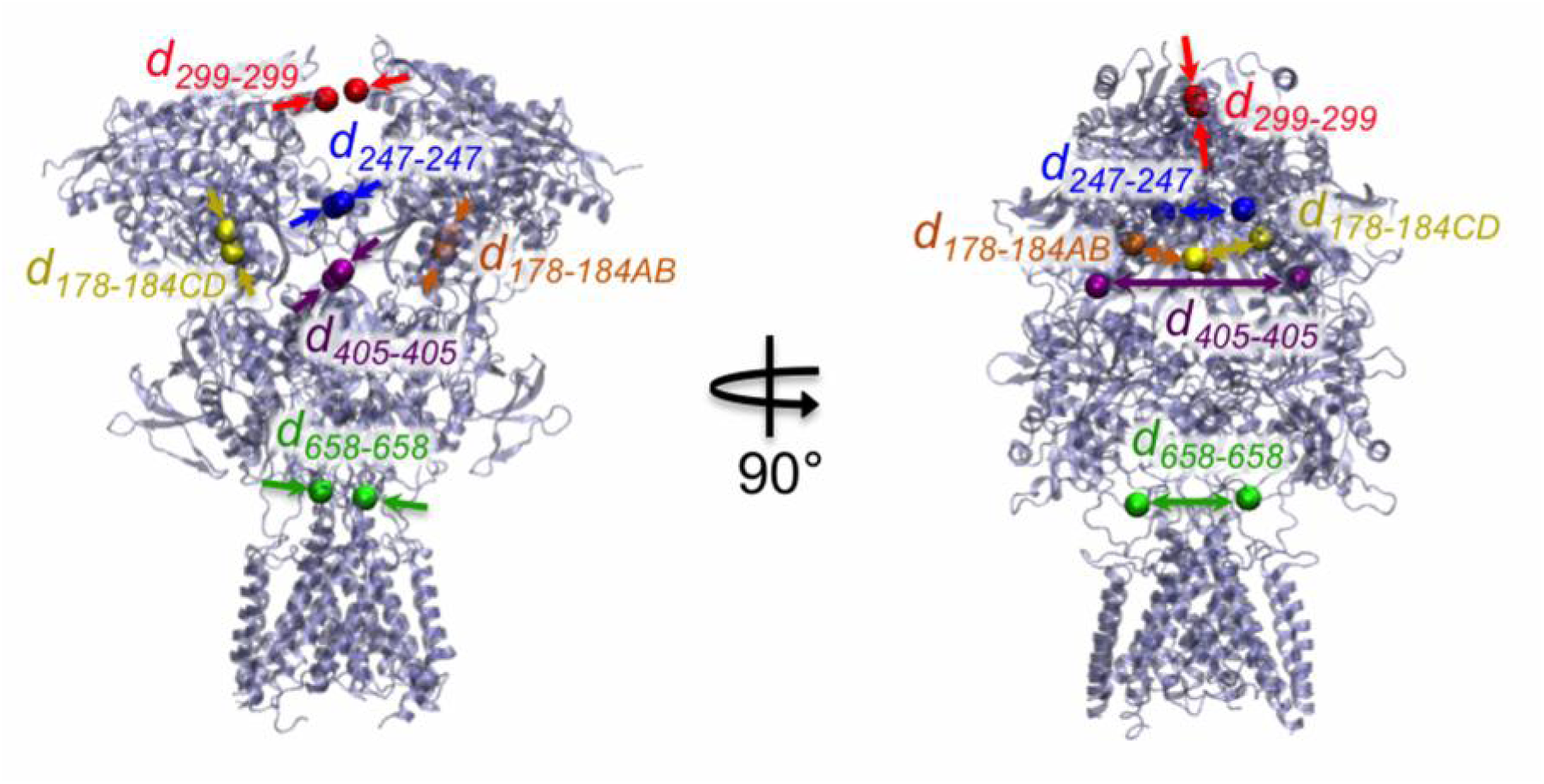
Characterization of a conformation of an NMDAR with six variables: *d*_*299-299*_ measuring the distance between the two heterodimers in the ATD, *d*_*178-184AB*_ and *d*_*178-184CD*_ measuring the rotation of the GluN1 and GluN2B subunits relative to each other within each of the two heterodimers, *d*_*247-247*_ measuring the rotation of the two heterodimers relative to each other in the ATD, and *d*_*405-405*_ and *d*_*658-658*_ measuring the rotation of two heterodimers in the LBD relative to each other.

Comparison of the intervals of the six distances sampled in our computations to those for the experimental structures shows that the simulations sampled a significant part, though not all of the conformational landscape (Table S2). In particular, all four ‘non-active’ cryoEM structures^7^ lie within the computationally sampled ranges. A more refined view on the relationship of the experimental structures to the sampled conformations, which takes into account correlations between these distances, is provided by the projection of the conformational landscape onto the first tICs (see below, section S4).

Some of the distances shown in Table S2 cannot be computed for the original PDB structures because of missing fragments in those structures. We made best possible estimates of these distances (shown in italics in Table S2) by partially reconstructing the corresponding structures. These values should be treated with caution. In particular, each of the *d*_*658-658*_ values for PDB structures 5IPQ, 5IPR, 5IPS, 5IPT, 5IPU where computed with the use of reconstructed coordinates of a C_α_ atom 9 amino acid residues away from the fragments present in the raw structure, and the same variable for PDB structure 5IPV was computed based on reconstructed coordinates of two C atoms, each 4 residues away from the fragments present in this PDB structure.

**Table S2.** The six characteristic distances in the computationally sampled conformations of NMDARs vs. those in the experimental structures, in Å. Four ‘non-active’ experimental structures are shown in bold. Estimates of missing data are shown in italics.

| | $d_{299-299}$ | $d_{178-184AB}$ | $d_{178-184CD}$ | $d_{247-247}$ | $d_{405-405}$ | $d_{658-658}$ |
| --- | --- | --- | --- | --- | --- | --- |
| Computationally sampled structures |  |  |  |  |  |  |
| holo | 3.7...44.5 | 11.1...22.0 | 8.2...22.0 | 3.9...32.2 | 39.2...60.9 | 16.9...36.3 |
| apo | 4.4...49.2 | 10.2...24.1 | 9.7...24.4 | 3.8...39.1 | 39.1...63.3 | 17.1...35.2 |
| Experimental structures |  |  |  |  |  |  |
| 4PE5 | 7.0 | 17.3 | 17.1 | 19.7 | 49.6 | 25.4 |
| 5FXG | 31.7 | 11.5 | 11.5 | 6.3 | 36.5 | 47.2 |
| **5FXH** | **22.6** | **17.1** | **17.1** | **21.3** | **53.1** | **27.9** |
| **5FXI** | **37.8** | **16.5** | **16.4** | **21.5** | **48.1** | **28.9** |
| **5FXJ** | **33.6** | **15.9** | **15.8** | **15.8** | **49.7** | **27.2** |
| **5FXK** | **37.1** | **16.3** | **16.3** | **19.0** | **48.4** | **29.1** |
| 4TLL | 11.9 | 17.1 | 16.9 | 13.2 | 51.9 | 25.1 |
| 4TLM | 11.8 | 17.2 | 16.1 | 12.7 | 52.1 | 26.9 |
| 5IOU | 35.6 | 15.9 | 13.3 | 5.8 | 50.3 | 28.8 |
| 5IOV | 23.8 | 17.4 | 16.6 | 14.6 | 51.5 | 29.1 |
| 5IPQ | 32.9 | 17.2 | 16.1 | 56.8 | 83.9 | *37.1* |
| 5IPR | 92.4 | 17.2 | 16.1 | 105.1 | 109.8 | *44.9* |
| 5IPS | 101.0 | 17.2 | 16.1 | 103.7 | 99.2 | *41.5* |
| 5IPT | 128.8 | 17.2 | 16.1 | 122.9 | *117.0* | *45.5* |
| 5IPU | 141.0 | 17.2 | 16.1 | 123.2 | 102.5 | *37.6* |
| 5IPV | 32.0 | 14.0 | 15.4 | 8.6 | 56.1 | *43.1* |
| 5UP2 | 23.3 | 15.7 | 16.5 | 12.8 | 47.8 | 27.4 |
| 5UOW | 29.6 | 15.2 | 12.6 | 9.3 | 48.5 | 27.6 |

Note also that PDB structures 4TLL, 4TLM, 5IOU, 5IOV, 5IPQ, 5IPR, 5IPS, 5IPT, 5IPU, 5IPV, 5UP2, 5UOW refer to NMDARs from a frog (*Xenopus laevis*),^6,8,9^ which differ from the human NMDAR not only by point mutations, but also insertions and deletions. Moreover, PDB structures 5UP2 and 5UOW refer to the GluN1/GluN2A/GluN2B triheteromeric NMDAR,^9^ while all other structures refer to the GluN1/GluN2B receptors. Therefore, a direct comparison of the interatomic distances for these 12 structures to the distances for our simulations and for the first 6 structures is to some degree questionable.

### S4. Physical interpretation of tICs

To find a simple physical interpretation of first tICs, it is useful to analyze pairwise correlations of tICs with the time series in the raw data, in this case, the six interatomic distances (Fig. S4, S5).

In the apo form, tIC 1 correlates mainly with *d*_*299-299*_ (*R*^2^=0.53) and *d*_*247-247*_ (*R*^2^=0.49), while correlation with the other four distances is much weaker (*R*^2^<0.05). tIC 2 closely correlates with *d*_*658-658*_ (*R*^2^=0.90), but not the other five distances (*R*^2^<0.13). tIC 3, like tIC 1, mainly correlates with *d*_*247-247*_ (*R*^2^=0.48) and *d*_*299-299*_ (*R*^2^=0.44), but not the other four distances (*R*^2^<0.13). Therefore, the slowest timescale dynamics revealed in our simulations of the apo form is related to movement of the two heterodimers in the ATD relative to each other (tIC 1,3) and, independently, dilation of the gating ring by motion of the LBD-TMD linkers (tIC 2). Interestingly, the distance between the LBD-TMD linkers is captured as a relevant slow degree of freedom by the model, even though no pore opening events associated with this motion were sampled in our simulations.

**Fig. S4.**
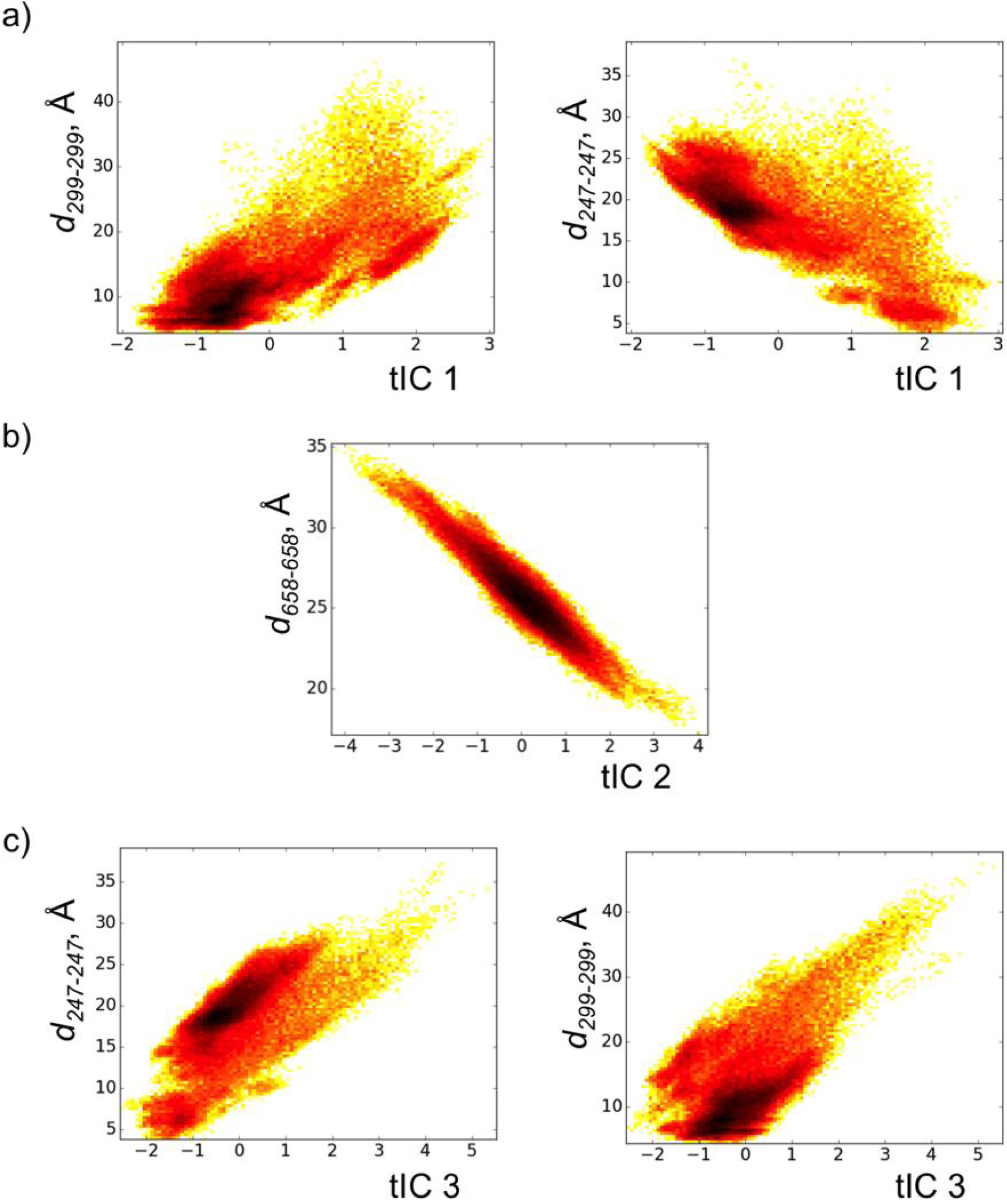
In the apo form, the first three tICs correlate mainly with the shown interatomic distances.

In the holo form, tIC 1 correlates mainly with *d*_*247-247*_ (*R*^2^=0.66) and *d*_*299-299*_ (*R*^2^=0.34), while correlation with the other four distances is weaker (*R*^2^<0.22). This collective variable is analogous to tIC 1 in the apo form, unlike the subsequent tICs. tIC 2 correlates with *d*_*178-184CD*_ (*R*^2^=0.64) and *d*_*405-405*_ (*R*^2^=0.34), but not the other distances (*R*^2^<0.09). tIC 3 mainly correlates with *d*_*658-658*_ (*R*^2^=0.55) and *d*_*299-299*_ (*R*^2^=0.27), but not the other four distances (*R*^2^<0.10). Therefore, the slowest timescale dynamics in the holo form, at least as sampled in our simulations, is different from the dynamics of the apo form. Besides the relative motion of the two ATD halves (tIC 1), it also includes the variables reflecting a coupling between motion of ATD-LBD linkers and changes in the relative position of GluN1 and GluN2B units within an ATD half (tIC 2), and coupling between the distance between the two ATD halves and that between the LBD-TMD linkers (tIC 3).

Automatic detection of the relevant degrees of freedom by tICA allows us to go beyond the analysis of individual interatomic distances, and account for their synchronized changes.

**Fig. S5.**
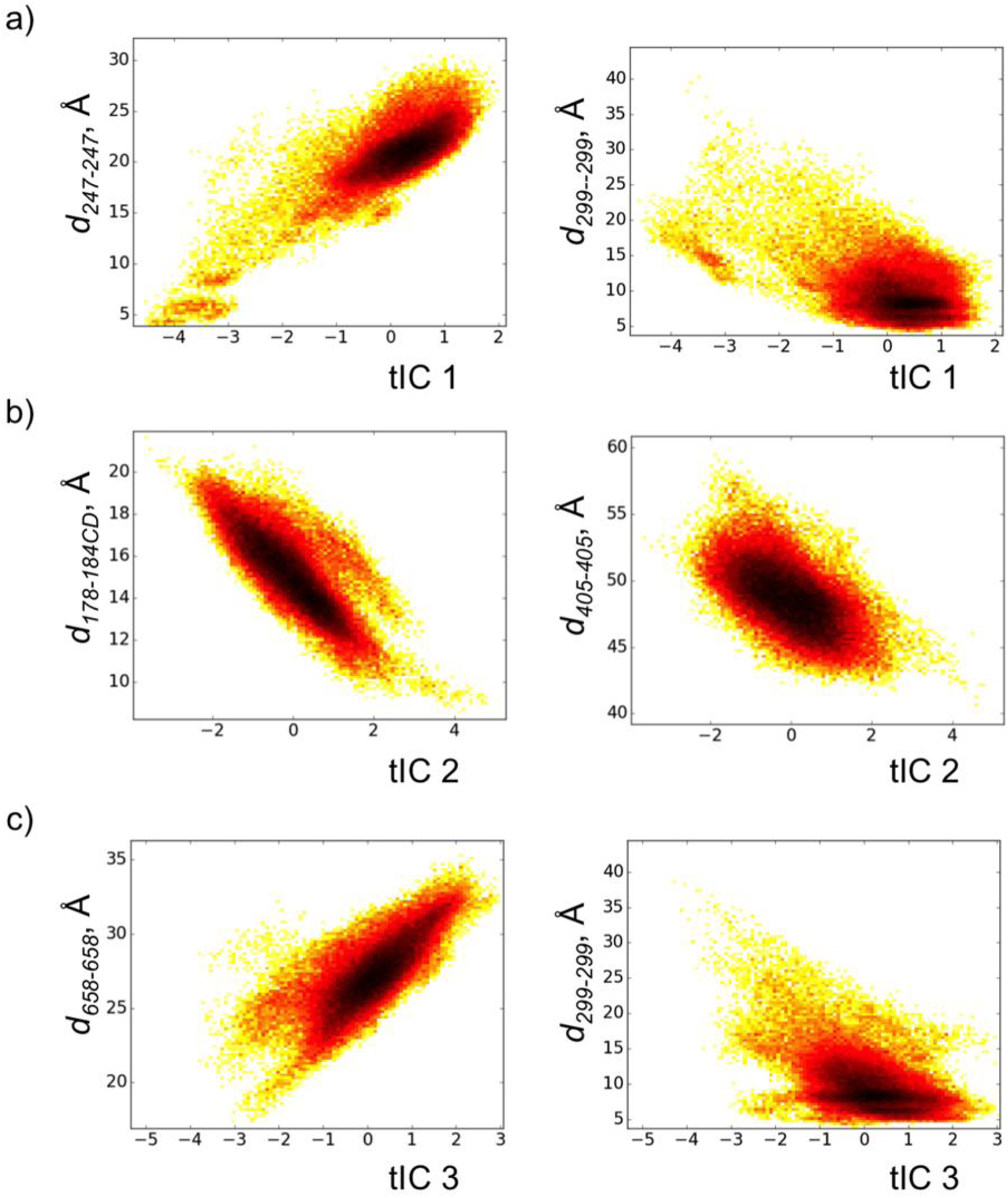
In the holo form, the first three tICs correlate mainly with the shown interatomic distances.

#### S5. Sampled conformation landscapes

Projections of the computationally sampled parts of the configuration space onto the subspaces spanned by the first three tICs are shown in Fig. S6 and S7. Fig. S6 shows the position of all currently available experimental structures in these subspaces. As mentioned in section S3, the interatomic distances and, therefore, tICs computed for PDB structures 5IPQ-5IPV, 5IOU and some others may be less reliable and less comparable to the simulated structures. Despite this uncertainty, it is evident that the cryoEM structures 5IPQ-5IPV (determined for antagonist-bound NMDARs) and 5IOU (the agonist-and-coagonist-bound ‘inactive’ state),^8^ as well as the ‘active’ cryoEM structure 5FXG,^7^ were not reached in the simulations. Presumably, these structures correspond to other functional states of the receptor different from the functional state sampled in our simulation, most probably, open or pre-open states. It is also interesting to note that the structure 5IPV is relatively close in these plots to the ‘active’ structure 5FXG, while the structures 5IPQ-5IPU are located in different parts of the tIC1/tIC2/tIC3 subspaces (sometimes in a direction opposite to 5FXG and 5IPV relative to the sampled region).

Fig. S7 shows only the experimental structures within or very close to the computationally sampled range. All the ‘non-active’ (shown in *green*), inhibited (*blue*), and some other structures (*yellow*, PDB codes 5IOV, 5UP2, and marginally 5UOW) were computationally sampled. The points corresponding to these experimental structures are scattered throughout the sampled region, though some of the sampled conformations do not have close analogues among the currently known experimental structures (for more details see sections S6.2-3).

**Fig. S6.**
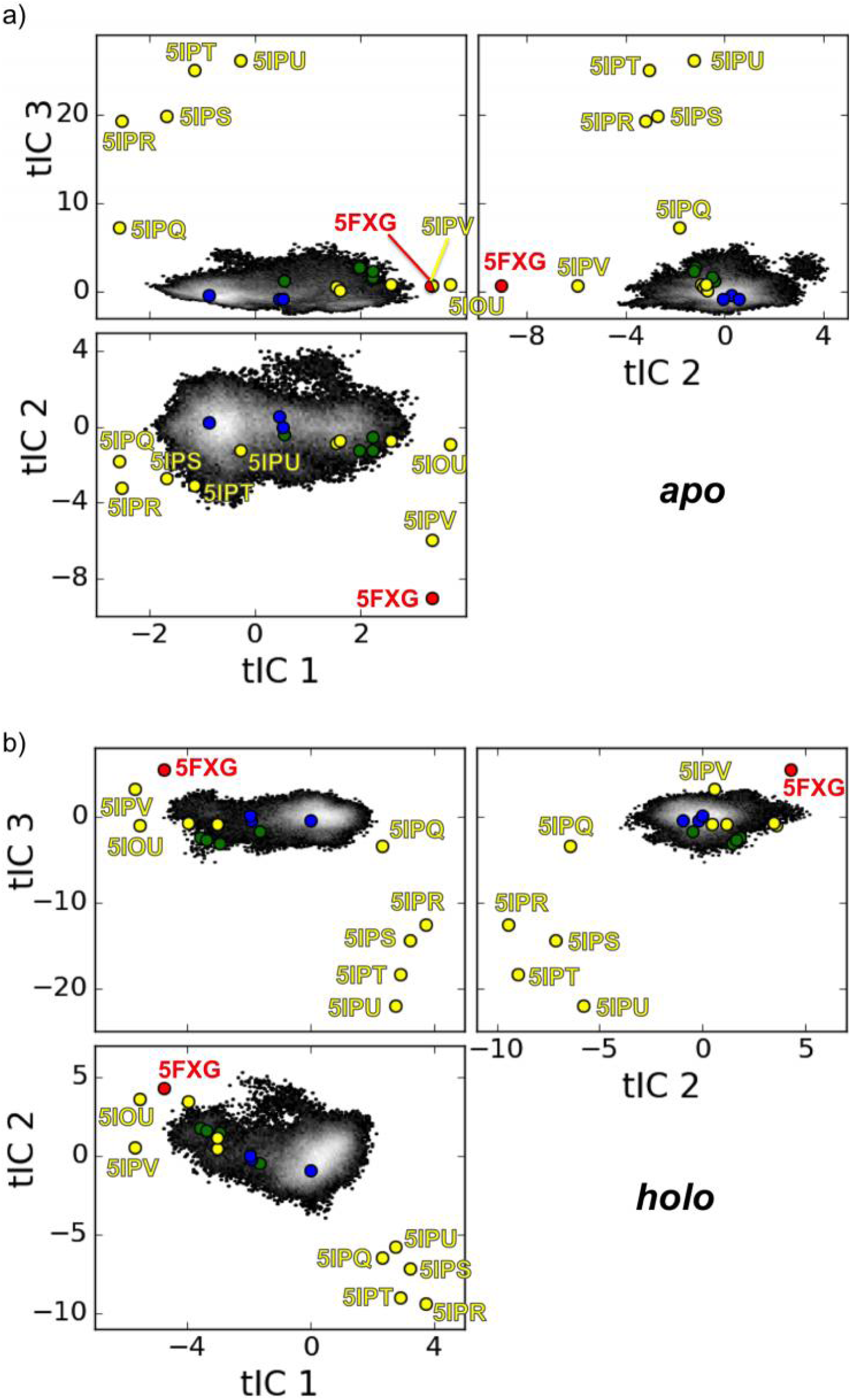
Conformation landscapes with the positions of all experimental structures. The color coding is the same as in Fig. 1b,c in the main text.

**Fig. S7.**
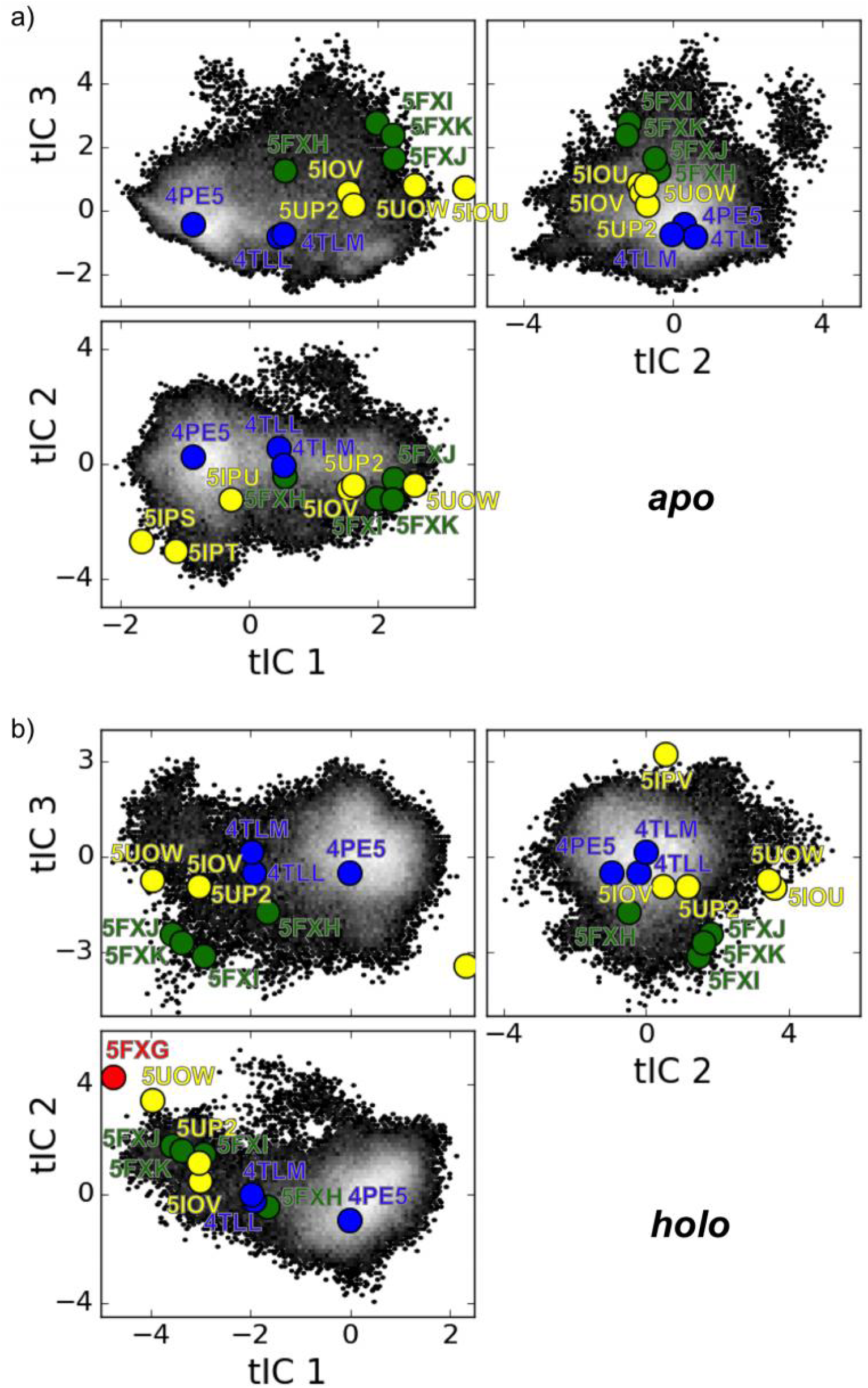
Conformation landscapes zoomed onto the computationally sampled region. The color coding is the same as in Fig. 1b,c in the main text.

### S6. Markov state models

#### S6.1. Choice of the number of Markov states

In the case of continuous distributions, it is not evident how many states to use in a Markov state model. An objective approach to this problem is provided by a cross-validation with the GMRQ scoring function.^36^

As expected, the performance of models steadily improves with the number of states, if it is scored against the same data that were used for fitting the models (Fig. S8, *blue*). However, the performance of the models estimated by cross-validation (*red*), when different subsets of raw data are used for training and testing, improves only to a certain number of states, and after that deteriorates due to overfitting. The maximal score computed with cross-validation defines the optimal number of states in the model.

For the simulations of the apo NMDAR, the optimal number of states turned out to be 197. Note that the maximum in the plot for the GMRQ score vs. the number of states is relatively shallow, showing that any number of clusters in the range between ∼150 and ∼300 would work nearly well. For the simulations of the holo NMDAR, the optimal number of states was 79. As in the previous case, the maximum in the plot for the GMRQ score vs. the number of clusters is rather wide, showing a small sensitivity of the quality of the model to the number of states in the range from ∼50 to ∼150. The difference between the optimal number of states for the generated sets of trajectories of apo and holo NMDARs is unclear to us and may simply be a result of a statistical fluctuation.

**Fig. S8.**
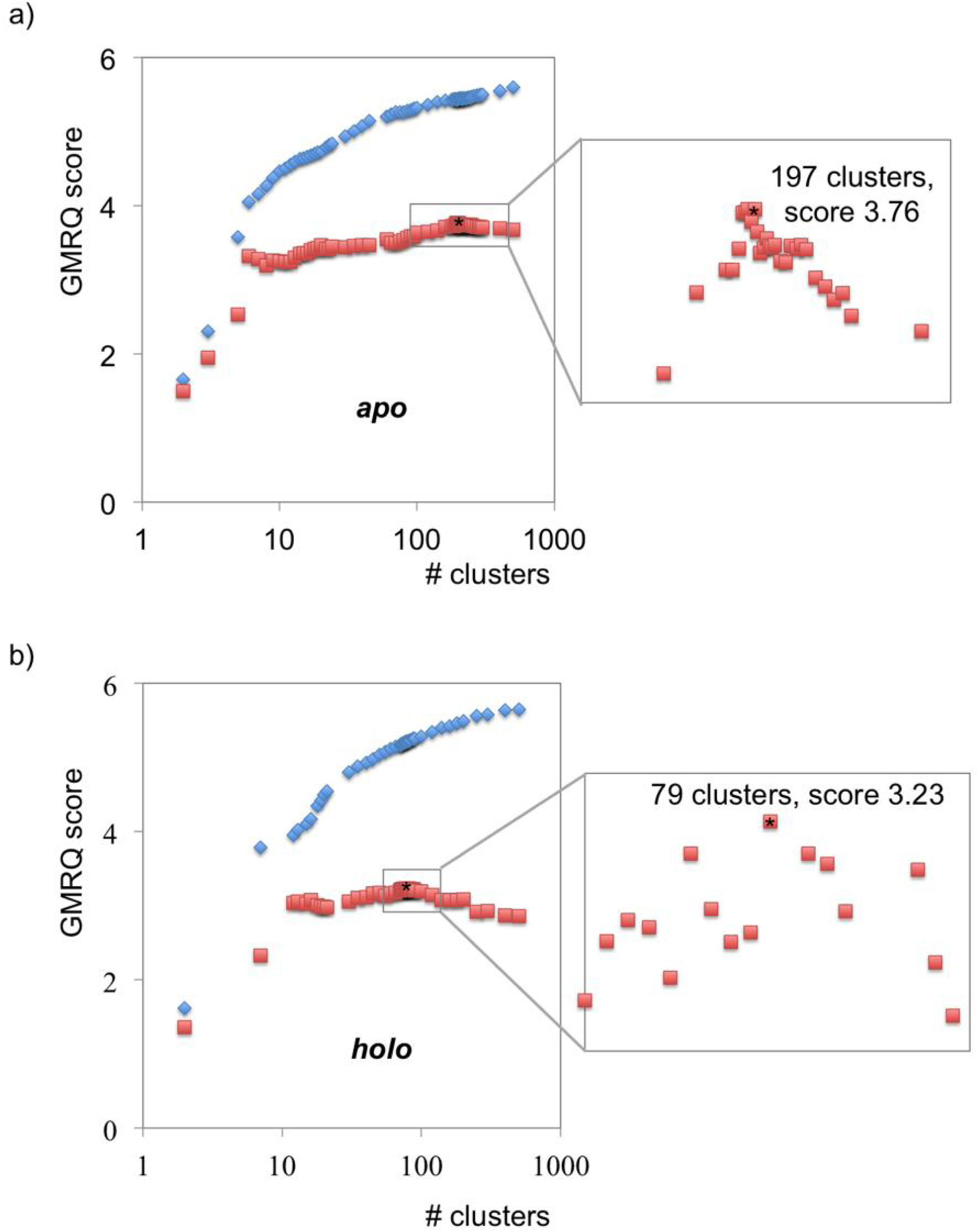
The number of states in the MSMs was chosen based on the GMRQ criterion.

#### S6.2. Timescales and their assignments

MSMs provide not only estimates of the characteristic timescales of processes in the system, but also their assignment in terms of specific states.^35^ In general, each timescale corresponds to a set of simultaneous transitions from all states to all other states with various weights determined by the corresponding eigenvector of the transition matrix. However, in practice such weights are far from zero for only a few states. In this case, the corresponding dynamic process can be interpreted as a transition from one (or a few) state with a well-defined geometry to another state (or a few states). It is convenient to make such assignments with the use of diagrams shown in Fig. S9, S10. In panels a-e in each diagram, the eigenvectors for the first five timescales are visualized as sets of circles. The color of each circle shows the value of the corresponding element of the eigenvector (heatmap coloring, large negative values shown with deep blue, large positive values with deep red), while the size of the circle is proportional to the absolute value of the corresponding element of the eigenvector reflecting its relative importance. Therefore, the process with the corresponding timescale is mainly a transition from the states shown with largest deep-red circles to the states shown with the largest deep-blue circles, or vice versa.

In the apo form, the longest timescale of 5.6 μs (Fig. 9a) corresponds to transitions from a diverse set of stable structures (including those close to the ‘non-active’ and inhibited structures) to a state that does not have close analogues among the experimental structures (denoted as state A in Fig. 9e). Timescale 2 (0.39 μs) corresponds to transitions from a diverse set of stable structures to another state without close analogues among the experimental structures (denoted as state B in Fig. 9e). Timescale 3 (0.29 μs) corresponds to transitions from inhibited geometries, especially 4PE5, to two different states. One of these two states is close to experimental structures 5IOV and 5UP2, and the other is the above-mentioned state B. Timescale 4 (0.16 μs) corresponds to transitions between the ‘non-active’ structures and inhibited structures 4TLL and 4TLM. Timescale 5 (0.15 μs) refers to transitions between the inhibited structures and one more state without close analogues among the experimental structures (denoted as state C in Fig. 9e).

In the holo form, the longest timescale of 0.99 μs (Fig. 10a) corresponds to transitions from a diverse set of stable structures (mainly inhibited) to a state that does not have close analogues among the experimental structures (denoted as state D in Fig. 10e). Timescale 2 (0.59 μs) corresponds to transitions from inhibited structures to another new state (denoted as state E in Fig. 10e). Timescale 3 (0.11 μs) corresponds to transitions between inhibited states, primarily 4PE5, and a diverse set of structures including the ‘non-active’ structures and structures with PDB codes 5IOV and 5UP2. Timescale 4 (0.10 μs) corresponds to transitions between inhibited states, particularly 4PE5, and states close to the experimental structure 5FXH (‘non-active 1’). Timescale 5 (0.08 μs) refers to transitions between inhibited states and state E.

Overall, our simulations have revealed a wealth of transitions on submicrosecond timescales (except for one multi-microsecond transition in the apo form on the timescale of 5.6 μs) between various conformations of NMDARs, including conformations close to several structures known from experiments (mainly, the ‘non-active’, inhibited and those with PDB codes 5IOV and 5UP2), as well as states that currently do not have close analogues among the experimental structures (denoted in this work as states A-E).

**Fig. S9.**
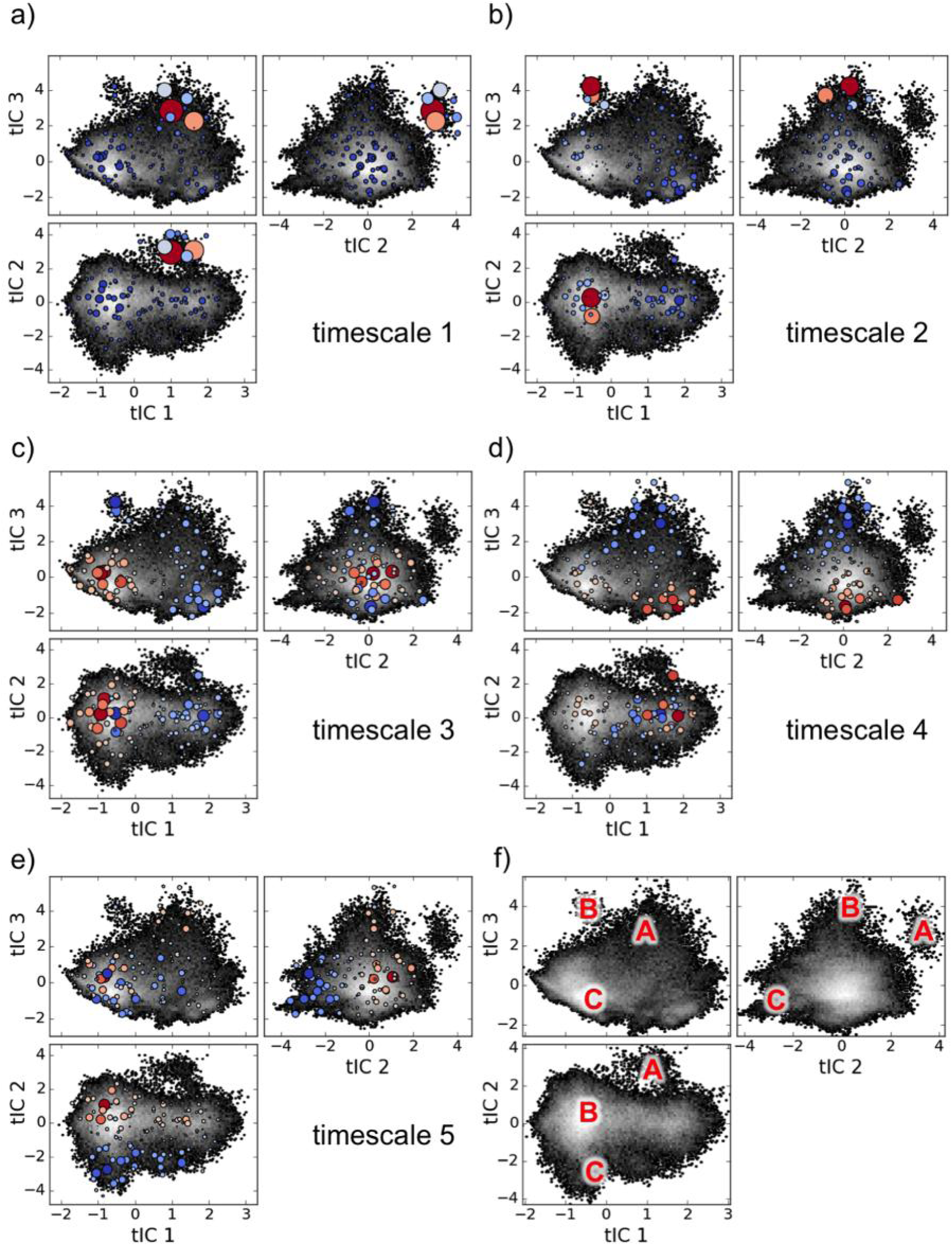
Assignment of the slowest timescale transitions in the apo form of NMDAR. (a-e) Transitions corresponding to the five slowest timescales. (f) Conformations of an apo NMDAR revealed by simulations without close analogues among the known experimental structures.

**Fig. S10.**
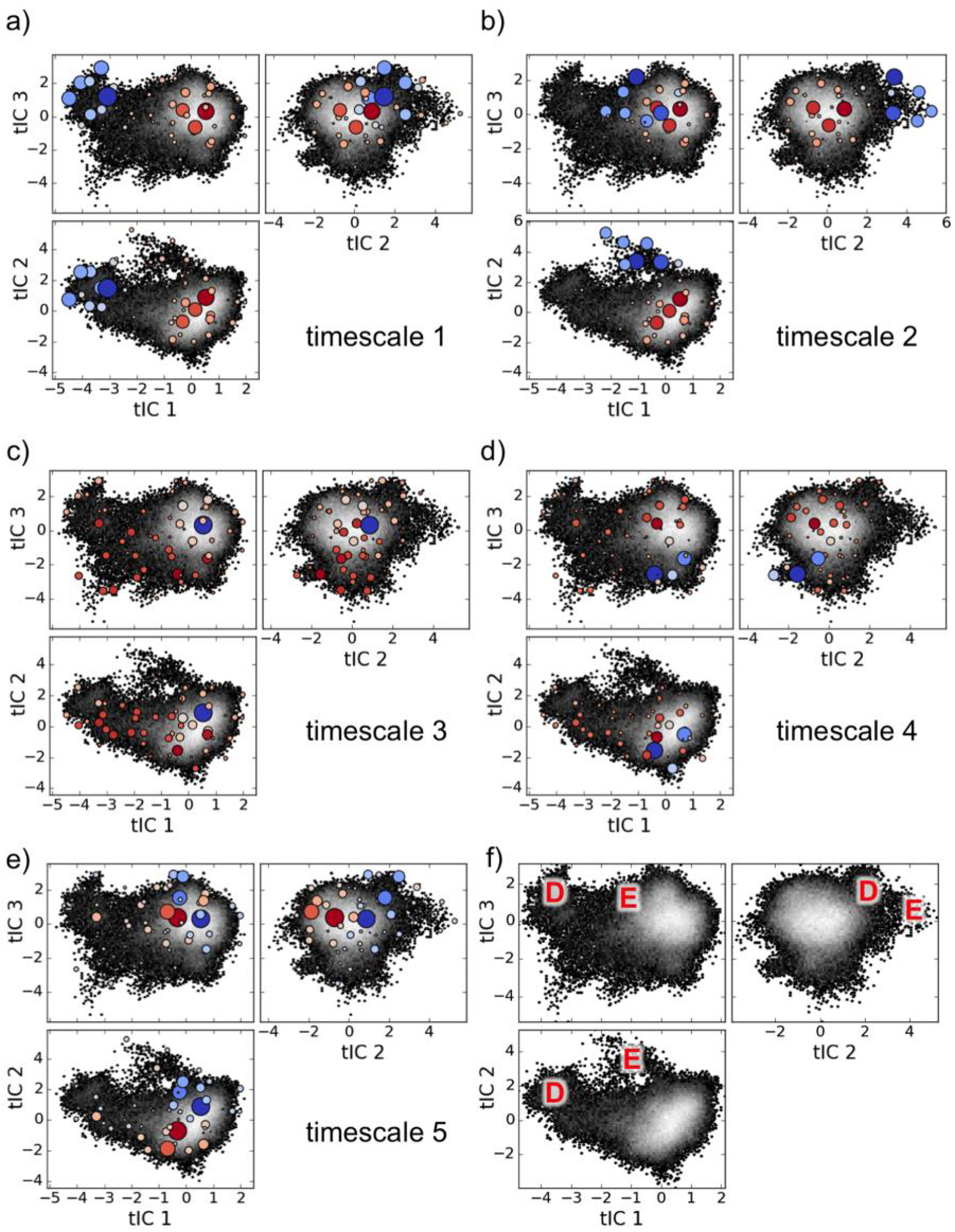
Assignment of the slowest timescale transitions in the holo form of NMDAR. (a-e) Transitions corresponding to the five slowest timescales. (f) Conformations of a holo NMDAR revealed by simulations without close analogues among the known experimental structures.

#### S6.3. Previously unknown predicted states of NMDAR

Our simulations have revealed five states that do not have close analogues among the currently known experimental structures (Fig. S9e and S10e). We expect that they may be biologically important because they were revealed by the analysis of the slowest timescale processes in the simulated apo and holo NMDARs. Below we briefly characterize these structures and their main difference from the experimental ones. tICs mentioned below refer to the apo form trajectories in the cases of states A, B and C, and the holo form trajectories in the cases of states D and E.

*State A* is characterized by large positive values of tIC 2, which correspond to small values of *d*_*658-658*_ (∼18 Å), and therefore a tightly closed gating ring at the boundary of the LBD and TMD, even tighter than in the inhibited structures 4PE5, 4TLL, 4TLM (*d*_*658-658*_ ∼25-27 Å). tIC 3 values for state A are also greater than the inhibited structures, implying that the distances between the ATD halves *d*_*299-299*_ and *d*_*247-247*_ are larger in this state, and therefore the ATD in state A is dissociated.

*State B* is characterized by large positive values of tIC 3, while the values of tIC 1 and tIC 2 are close to 0. In this state, the distances between the ATD halves *d*_*299-299*_ and *d*_*247-247*_ are larger than in all other states, experimental and predicted here, and therefore the ATD in state B is widely dissociated (*d*_*299-299*_ ∼30-45 Å, *d*_*247-247*_ ∼25-35 Å). The difference from state A is that the gating ring in state B does not close so tightly (*d*_*658-658*_ ∼25 Å). The difference from the experimental ‘active’ geometry (5FXG) is in the relative rotation of the ATD halves. In 5FXG, they are rotated such that the distance between residues 247 is relatively small (*d*_*247-247*_ = 6.3 Å), while in state B their angle of rotation is different, and this distance reaches ∼25-35 Å. At the same time, the distances between the ATD halves (measured by *d*_*299-299*_) in state B and in 5FXG are comparable.

*State C* has large negative values of tIC 2, corresponding to a relatively dilated gating ring. The distance between the LBD-TMD linkers *d*_*658-658*_ in this case is ∼35 Å, which is much larger than in the inhibited and ‘non-active’ experimental structures (25-29 Å), but less than in the ‘active’ structure 5FXG (47 Å) or the structure 5IPV (43 Å).

*State D* corresponds to large negative values of tIC 1, and therefore small values of *d*_*247-247*_ (∼5 Å) (as seen from Fig. S5a, large distances are not achieved for such extreme values of tIC 1, despite the overall negative slope in the pairwise correlation). These values correspond to a mutual orientation of the ATD halves where they nearly touch each other, but in a rotated position. Among the experimental structures, similar arrangement is observed in PDB entries 5IOU (the agonist-and-coagonist-bound ‘inactive’ state) and 5FXG (the agonist-and-coagonist-bound ‘active’ state). However, the structure 5FXG has a much wider gating ring (47 Å, in comparison to ∼30 Å in state D and 29 Å in 5IOU). A comparison of the landscapes given in Fig. S10e and S6b shows that state D is ∼2/3 the way between the initial geometry and 5IOU, implying that state D might merge with the structure 5IOU with more computational sampling.

*State E* is characterized by large positive values of tIC 2, and therefore small values of *d*_*178-184CD*_ (∼9 Å) and *d*_*405-405*_ (∼43 Å). This corresponds to relative rotation of GluN1 and GluN2B subunits within one of the ATD halves and a decrease in the distance between two ATD-LBD linkers, respectively. In both regards, this state resembles the ‘active’ cryoEM structure (except that in the ‘active’ structure the relative rotation of GluN1 and GluN2B subunits occurs in both ATD halves, while here in only one of them). However, state E is different from the ‘active’ structure in the tIC 1 values: for state E, tIC 1 is closer to 0 (implying a mutual arrangement of the ATD halves like in 4PE5), while for the ‘active’ state tIC 1 = –4.8 (implying a large distance between residues 299 in the ATD halves, but a small distance between residues 247).

To sum up, the computationally predicted states A-C, E look like chimeric structures joining various features from different known experimental structures, while state D may correspond to the experimental state 5IOU (the agonist-and-coagonist-bound ‘inactive’ state) not completely reached due to limited sampling. In all of these cases, the predicted structures look reasonable and may serve, for example, to guide future experiments (e.g., by motivating optimal locations of spectroscopic probes on NMDARs) or for *in silico* drug design.

### S7. Description of the attached videos

#### Video S1

Simulated molecular systems included the protein part of the NMDA receptor, two molecules of an agonist (glutamate) and two molecules of the coagonist (D-serine) (only in the holo form), glycans covalently attached to the protein, a lipid bilayer, and a water solution of NaCl (see section S2).

A lower-resolution version of Video S1 is publically available at: https://youtu.be/6uVmIDsXyDA

#### Video S2

An MD trajectory of an NMDAR (apo form, trj 124) showing transitions to the ‘non-active’ states in less than 100 ns. For notations and color coding, see Fig. 1b,c and 2a from the main text.

A lower-resolution version of Video S2 is publically available at: https://youtu.be/aJsNE0zL32s

#### Video S3

Another MD trajectory of NMDAR (apo form, trj 26) that started from the same initial geometry as the trajectory in Video S2, but did not exhibit an ATD dissociation.

A lower-resolution version of Video S3 is publically available at: https://youtu.be/CLnMf_Y1fEM

